# Cryptic host phenotypic heterogeneity drives diversification of bacteriophage λ

**DOI:** 10.1101/2024.08.05.606710

**Authors:** Caesar A. De La Fuente, Nehme Lahoud, Justin R. Meyer

## Abstract

Bacteriophages, the most abundant and genetically diverse life forms, seemingly defy fundamental ecological theory by exhibiting greater diversity than their numerous bacterial prey. This paradox raises questions about the mechanisms underlying parasite diversity. To investigate this, we took advantage of a surprising experimental result: when bacteriophage λ is continually supplied a single host, λ repeatedly evolves multiple genotypes within the same flask that vary in their receptor use. Measurements of negative frequency-dependent selection between receptor specialists revealed that diversifying selection drove their evolution and maintenance. However, the source of environmental heterogeneity necessary to generate this type of selection was unclear, as only a single isogenic host was provided and replenished every eight hours. Our experiments showed that selection for different specialist phages oscillated over the 8-hour incubation period, mirroring oscillations in gene expression of λ’s two receptors (*Escherichia coli* outer membrane proteins LamB and OmpF). These receptor expression changes were attributed to both cell-to-cell variation in receptor expression and rapid bacterial evolution, which we documented using phenotypic resistance assays and population genome sequencing. Our findings suggest that cryptic phenotypic variation in hosts, arising from non-genetic phenotypic heterogeneity and rapid evolution, may play a key role in driving viral diversity.

## Introduction

Phages, the viruses that infect bacteria, exhibit an extraordinary degree of genetic diversity, and are estimated to be the most abundant life form on earth with 10^31^ virions globally^1,2^. This remarkable abundance and diversity are essential for numerous functions of microbial communities, including the health-benefits provided by the human microbiomes^3,4^ and marine biogeochemical cycling^5^. However, the underlying mechanisms driving the evolution of such high levels of phage diversity remain poorly understood.

Longstanding niche theory posits that resource heterogeneity promotes biological diversification^6,7^. In the context of phages, the diversity of their resources – bacterial species – offers many opportunities for niche creation given high bacterial species diversity^8,9^. Nevertheless, phage genetic diversity often surpasses bacterial diversity across divergent habitats,^10,11^ and has even been recorded as possessing 10 times higher species dievrsity^12,13^. This imbalance between phage and bacterial diversity contradicts the predictions of food-chain models, which posit that predator diversity should collapse when it outnumbers prey diversity^14-16^. Furthermore, theoretical models of phage-bacterial coevolution predict the one-to-one correspondence between the evolution of phage diversity and host diversity^17^.

The observed imbalance between phage and host diversity suggests that mechanisms beyond host genetic diversity must be driving and maintaining phage diversity. Additional evidence comes from the fact that bacteria can be infected by multiple phage species^18-20^. Closely related phages that infect the same host have evolved mechanisms to maintain species boundaries in the face of coinfection and hybridization^21^, suggesting there are evolutionary forces at play maintaining phage diversity on single hosts. Collectively, these findings imply that phages must partition their environments more finely than at the host-strain level. Direct empirical tests of the requirement for host genetic diversity in the evolution and coexistence of multiple phage species are still lacking. This knowledge gap has significant implications for our understanding of biodiversity and the evolution of species diversity.

Recently, we made a discovery while evolving bacteriophage λ in the laboratory that may help explain how phage diversity can outstrip bacterial diversity. In a previous study, a laboratory-evolved λ (EvoC) that gained use of a second receptor (OmpF, LamB is its native receptor)^22^ was shown to diversify into two receptor specialists when provided with two hosts, each expressing either OmpF or LamB^23,24^. We repeated this study, but with a control group consisting of a single host that expresses both receptors (wild type, ‘WT’). We predicted that EvoC would not diversify in this control group, as there should be minimal host heterogeneity. Instead, we expected a single lineage adapted to WT to sweep through the population, eliminating genetic variation. However, EvoC repeatedly diversified into a range of receptor generalists and specialists, resulting in genetically diverse populations of phage being maintained on a single bacterial host. This counterintuitive observation mirrors the imbalanced ratio of phage-to-hosts in nature, prompting us to conduct a series of studies on this failed control.

Several hypotheses may explain why λ diversity evolved despite the presence of a single bacterial host. One possibility is that the changes are selectively neutral, as all host cells possess the capacity to express both receptors, allowing receptor tropism to drift freely without affecting phage fitness. However, this neutral evolution hypothesis seems less likely given the short duration of the experiment (280 hours) and the large phage population sizes (averaged 10^7^-10^8^ virions). Alternatively, phage diversity could evolve due to natural selection if there was heterogeneity among the cells, effectively creating a multi-host condition that promotes diversification. Gene expression variability could arise from random fluctuations in cell-to-cell expression, but the cells were in the exponential phase of growth, which tends to reduce gene expression variability^25^. Receptor heterogeneity may also arise if the cells adapt to repress expression of one or the other receptor to protect from infection^26,27^; however, WT hosts were resupplied every eight hours, and this period was thought to be too short for resistance mutants to evolve to appreciable frequencies.

To test these hypotheses, we performed a series of experiments on evolving phage populations. By sequencing the phage’s host-recognition gene (*J*), we detected nonrandom patterns of substitution consistent with adaptive evolution, thereby excluding neutral evolution as a driver of phage diversity. We further demonstrated that phage tropism experienced natural selection, with a diversifying mode of selection driven by temporal oscillations in selection for divergent phage receptor tropisms. Notably, these oscillations in selection were closely coupled to oscillations in receptor gene expression, which were attributed only partly to rapid resistance evolution and must have also arisen due to non-genetic cell-to-cell phenotypic variability. Our results provide strong evidence for the hypothesis that bacterial phenotypic heterogeneity, arising from both cell-to-cell variability and rapid evolution, can drive phage diversification.

## Results

### Repeated λ diversification on a single host

In line with a previous study^23^, we propagated six populations of EvoC for 35 eight-hour growth cycles, but this time using a single host, WT, that expresses both receptors. After each cycle, phage particles were removed from the bacteria, and 1% of the population was transferred to a new flask of medium and a new supply of WT cells growing exponentially. Every 5th cycle, a phage sample was cryopreserved, and on the 25th, 30th, and 35th cycles four phages were isolated from each population: two from plaques that formed on bacterial lawns expressing OmpF and two from lawns expressing LamB. We measured the receptor tropism of the phages using efficiency of plaquing assays on LamB^-^ or OmpF^-^ host lawns^23^. Specialization indexes were computed to compare phage receptor preferences [Specialization index = (density on OmpF^-^ – density on LamB^-^) / (density on OmpF^-^ + density on LamB^-^)}], where values range from +1 (complete LamB specialization) to –1 (complete OmpF specialization).

EvoC exhibited repeated evolution of diverse receptor preferences across all six populations (Fig. 1). Notably, 15 of the 69 isolates displayed extreme receptor preferences, with 11 isolates exhibiting a strong bias towards LamB (+1) and 4 isolates towards OmpF (-1). Furthermore, 13 isolates demonstrated a pronounced OmpF-bias, while 28 isolates favored LamB, and only 13 isolates exhibited generalist behavior (receptor preference values between -0.33 and +0.33). The number of isolates that favor LamB is consistent with previous findings that EvoC more readily evolves to specialize on LamB than OmpF^23^. Importantly, every population exhibited variation in receptor tropism at least once during the experiment, although this variation shifted over time and diverged between populations. The observed stochastic patterns of diversity are consistent with the hypothesis that phage tropism evolution occurred via neutral drift, although the time scales should be too short to observe such significant drift and fluctuations are also consistent with shifting selection regimes.

**Fig. 1:**
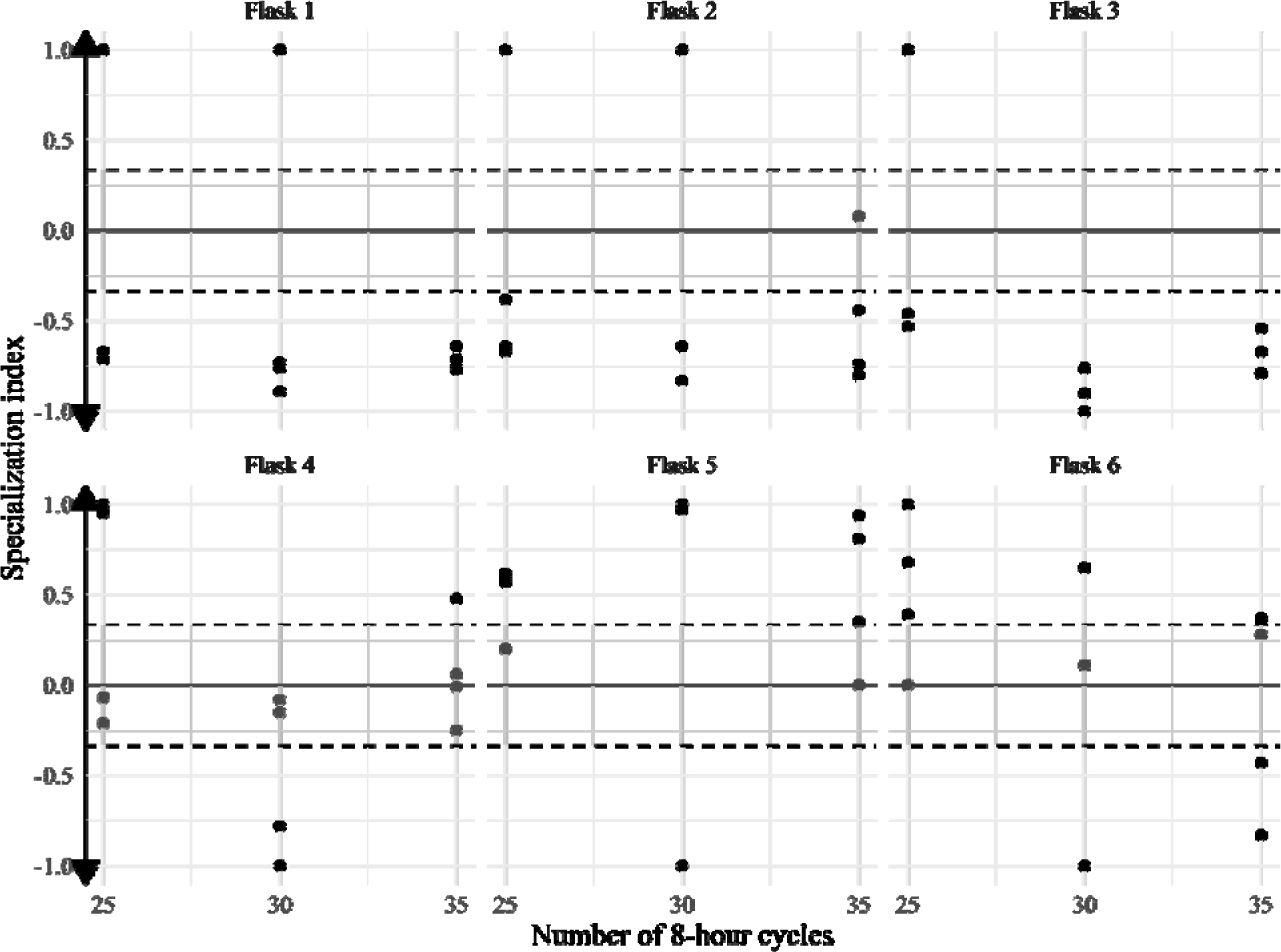
Specialization indices of phage λ isolates during the last 10 cycles. Specialization index ranges between +1 and –1, indicating complete LamB specialization to complete OmpF specialization. Grey area outlined by dashed lines indicate +/- 0.33 values and a transition from generalist to specialist as phages gain strong preferences. Flasks 3, 5, and 6 are missing one value each on day 30 due to plating errors.

### Nonrandom patterns in DNA sequence variation suggests adaptative evolution

Previous research has demonstrated that mutations within the reactive region of the phage receptor-binding protein (J) are responsible for alterations in phage receptor utilization^28^. To investigate whether similar evolutionary patterns occurred in this study, we sequenced the receptor-binding domain of the *J* gene. We selected eight isolates from cycle 30 of experimental flasks 2 and 5, as these samples exhibited the most pronounced intra-flask divergence in receptor tropism. Our analysis revealed 21 nonsynonymous mutations out of 23 single-nucleotide polymorphisms, suggesting that most of these changes were driven by natural selection (Fig. S1). Notably, all nonsynonymous mutations evolved in parallel across both flasks, a pattern consistent with natural selection rather than genetic drift^22^. For instance, all isolates acquired a C-to-T mutation at J gene nucleotide position 3310, while nearly all had a C-to-A mutation at position 3033. Furthermore, all λL phages (isolated on OmpF^-^ cells) gained a C-to-A mutation at position 3377, implying that this mutation may be responsible for LamB specialization and represents a signature of ecological divergence at the DNA sequence level^23^. Collectively, these findings suggest that natural selection, rather than genetic drift, drove the evolution of receptor tropism in our study.

### Negative frequency-dependent selection drives phage receptor-use variation

We investigated whether negative frequency-dependent selection occurs between receptor-specialist phages, a mechanism that favors rare phenotypes over common ones, thereby maintaining population diversity^29^. This type of selection can emerge from environmental niche heterogeneity, when one niche becomes overutilized by a population, selection shifts to favor rare genotypes^30,31^. Based on these principles, we hypothesized that the frequency of receptor-infecting phenotypes in the population would influence the direction of selection. Specifically, we predicted that artificially increasing the frequency of one receptor-specialist would lead to niche saturation, reducing its overall fitness, while the phage infecting the less-utilized receptor would benefit from being below its niche’s capacity, resulting in positive fitness.

To test the hypothesis of negative frequency-dependent selection, we conducted a competitive interaction experiment using two λ isolates from population #2, λL (adapted to use LamB) and λF (adapted to use OmpF). We genetically marked each phage with lacZα^32,33^, allowing us to distinguish plaques on bacterial overlay indicator plates. The genetically marked phages produce bright blue plaques and are easier to count when rare relative to typical clear plaques, and so when we set up competitions the genetically marked variant was always rare. Three replicates were set up with λF-lacZα rare and λL common and three with λL-lacZα and λF common. This was repeated three separate times (Trials A-C). We quantified the frequency of each genotype at the beginning and end of the 8-hour competition experiment and calculated λF’s relative fitness to λL. Our results consistently showed negative frequency-dependent selection across all three trials (trial A: adjusted R² = 0.91, p = 0.0016; trial B: adjusted R² = 0.95, p = 0.0004; trial C: adjusted R² = 0.86, p = 0.0048; Fig. 2). Notably, the frequency-dependent fitness lines intersected the x-axis between 0 and 1 in each trial, indicating a frequency at which the variants are predicted to exhibit equal fitness and could coexist. These findings support our prediction that, despite efforts to create environmental homogeneity, our experimental protocol inadvertently introduced sufficient heterogeneity to select for diverse phage populations, specifically those with divergent receptor preferences.

**Figure 2:**
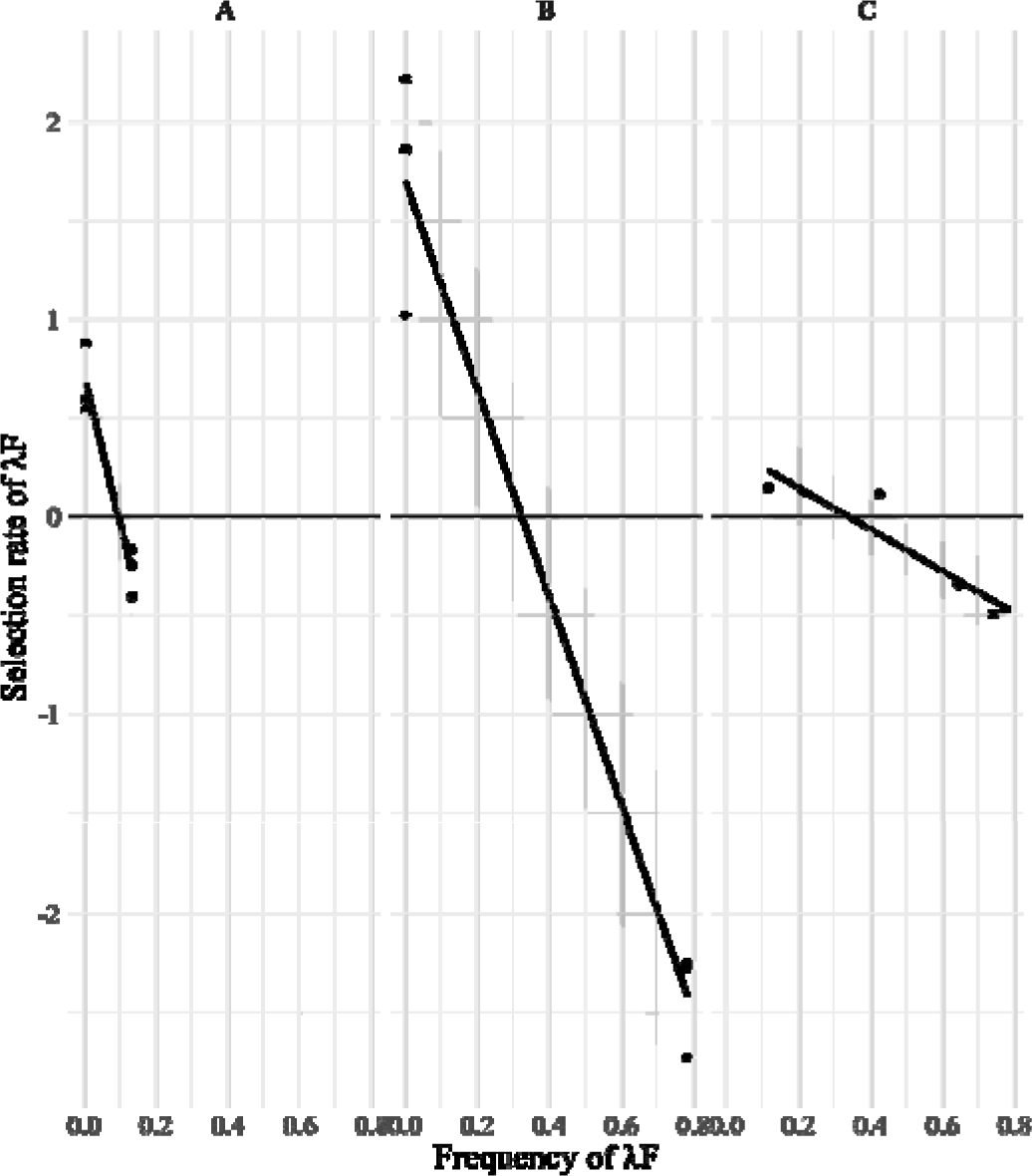
Frequency-dependent fitness functions measured in three separate trials. Selection rate is the difference in Malthusian growth parameters divided by time. Grey regions indicate 95% confidence intervals. Each trial showed the same pattern, although the strength of the frequency-dependence and the position of the fitness equilibrium point (the frequency where the line crosses the x-axis) varied by trial.

### Selection for receptor specialists fluctuates

Negative frequency-dependent selection can generate fluctuating selective pressures, where the dominant ecomorph is selected against, leading to a decrease in its frequency, and subsequently, selection flips to favor the previously disfavored type once it becomes rare. The shift in selection can be delayed, resulting in slight overshooting of the equilibrium frequency and course correction, causing temporal shifts in selection that oscillate back and forth with diminishing amplitude over time^34^. To further investigate diversifying selection in our system, we repeated the competition experiments, this time recording temporal dynamics by measuring selection every two hours. We conducted two trials, the first where λF was initially at a numerical advantage (selective disadvantage), and a second where λL was at a numerical advantage. Our results support our prediction: selection fluctuate over time (Fig. 3). In the first trial, λF initially exhibited a fitness disadvantage (average selection rate = -2.41 ± 0.27), but then became more fit than λL (average selection rate = 1.54 ± 0.09; glht p-value < 0.001). Subsequently, λF’s fitness decreased in each time interval (average selection rates = 0.39 ± 0.19 and -1.26 ± 0.38; glht p-values = 0.00151 and <0.001, respectively). Conversely, when λL was initially at a numerical advantage, the opposite dynamic emerged, with the last two intervals approached equilibrium between specialists (average selection rates = 0.69 ± 0.05, -0.73 ± 0.10, -0.10 ± 0.06, and -0.02 ± 0.05; glht p-values < 0.001, <0.001, and 0.338, respectively). These findings suggest that fluctuating selection helps drive the evolution and maintenance of phage genotypic diversity.

**Figure 3:**
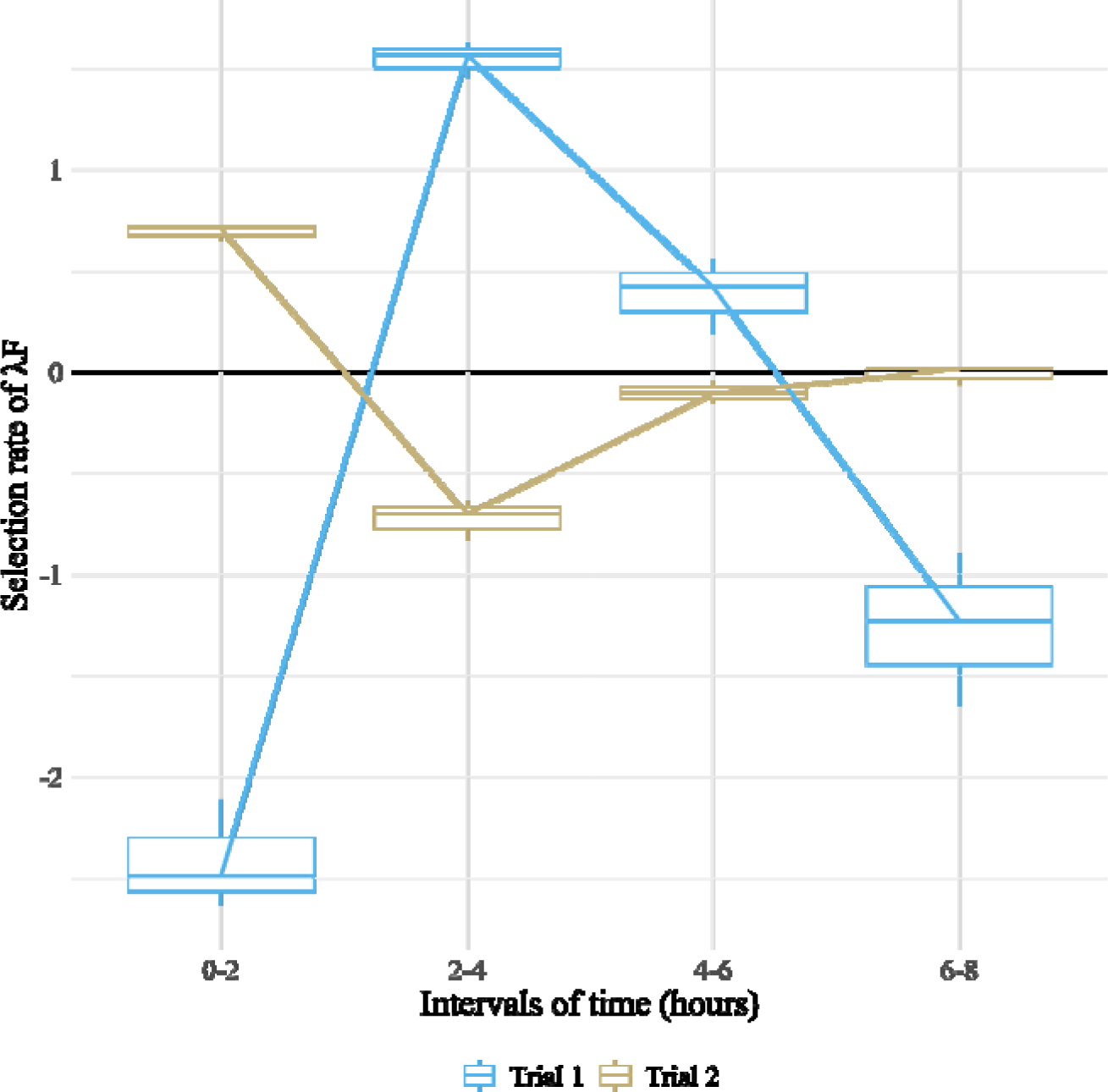
Selection fluctuates as genotypic frequencies equilibrate. Trial 1 was initiated with more λF than λL (λF averaged 72.15%). Trial 2 was run identically but initiated with more λL than λF (λL averaged 82.57%). Selection rate was measured over 2-hour intervals for the full 8 hours. Selection rate is the difference in Malthusian growth parameters divided by time. Boxplots display data from three replicates and convey the median, quartiles, and extreme maximum and minimum values.

### Receptor expression shifts within hours

After observing rapid fluctuations in selection for λF and λL, we hypothesized that these shifts were driven by changes in population-level expression of LamB and OmpF. We predicted that periods of higher OmpF expression would favor λF selection, while periods of higher LamB expression would favor λL selection. This would in turn drive oscillation in λF and λL frequencies. To test this, we repeated the temporal-varying selection experiment, monitoring LamB and OmpF gene expression using quantitative PCR. Our results showed that expressions of LamB and OmpF fluctuated throughout the experiment [ANOVA of LamB expression across time p-value = 7.7e-05 with F-value = 20.84; ANOVA of OmpF expression across time p-value = 6.06e-10 with F-value = 246.6] and that these fluctuations correlated with phage genotypic frequencies (Cross Correlation Tests; LamB with λL: corr = -0.995, p = 0.0003, no lag; OmpF with λF: corr = 0.996, p = 0.0033, 2-hour lag). The cell population initially showed much higher expression levels of OmpF relative to LamB, leading to a surge in the λF population by the second hour due to ample receptor availability. As λF frequency increased, OmpF expression dropped, as did λF frequency (Fig. 4). LamB started with low expression relative to OmpF, and λL initially lost ground to λF. By the second hour, the bacterial population increased LamB expression, allowing λL to regain prominence by the 4th hour, after which LamB expression dropped. Despite starting each 8-hour trial with an isogenic population of exponentially growing host cells, we observed substantial shifts in receptor expression, accounting for the changes in phage frequency and selection. These fluctuations provide one explanation for the negative frequency-dependent selection we measured between the receptor-specialists and play a role in phage diversification.

**Figure 4:**
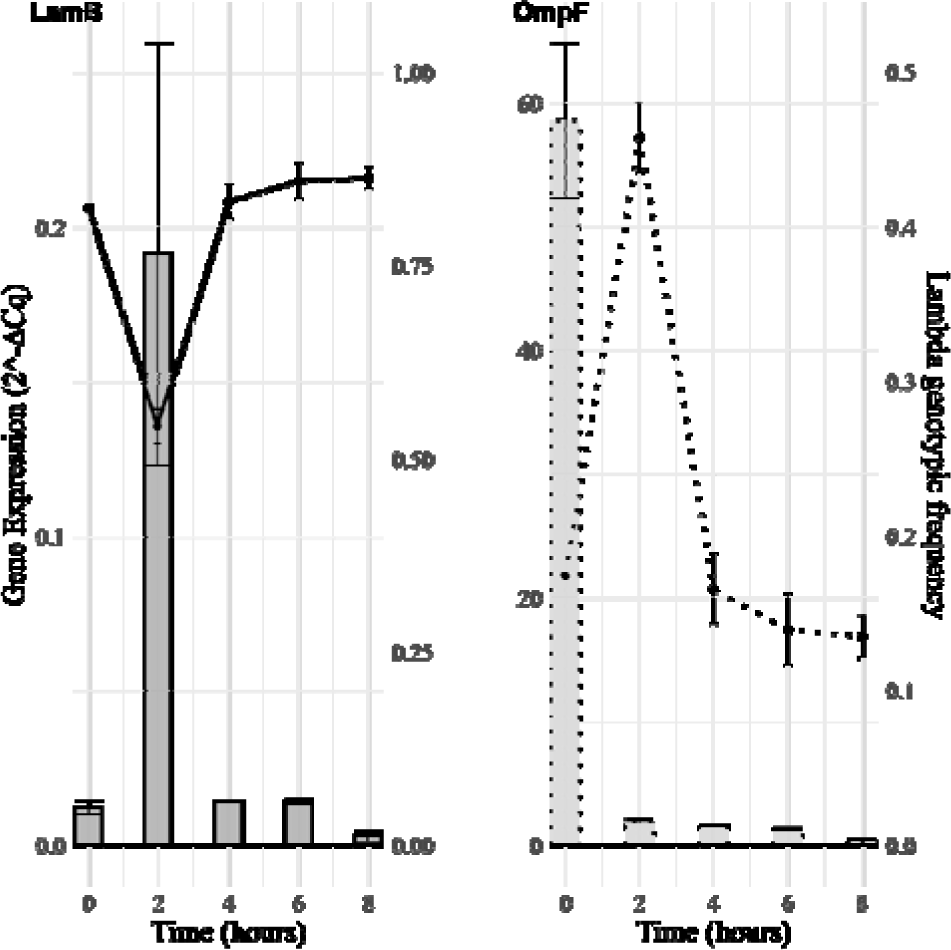
LamB and OmpF gene expression varied over time (bars). Expression levels were normalized to the constitutively expressed housekeeping gene, gapA. Error bars represent standard deviations, with each bar representing the average of triplicate measurements. Lambda genotypic frequencies (lines) fluctuated in response to their preferred receptor. Points represent the average of three replicates, with error bars indicating standard deviation. Notably, LamB expression and λL are perfectly out of phase, while λF’s frequency lags OmpF expression by 2 hours. Although the coarseness of the time series may contribute to this difference, the significant synchronization suggests feedback between receptor expression, selection, and phage genotypic frequencies.

### Testing the cause of receptor expression shifts

There are two main hypotheses to explain the observed shifts in gene expression: 1) non-genetic phenotypic heterogeneity and 2) genetic changes caused by rapid evolution.

The first hypothesis posits that temporal shifts in gene expression could be due to cell-to-cell variation. Bacterial cells are known to exhibit variation in gene expression even within the same population^35^, and this non-genetic variation can influence their sensitivity to phages^27^. It’s possible that there is phenotypic variation among cells in the expression of LamB and OmpF, and population-wide expression could fluctuate as different phage types rise to prominence and disproportionately infect cells that over-express their preferred receptor. Additionally, temporal variation in gene expression may occur over the 8-hour period due to environmental changes that alter gene expression, such as shifts in resources (sugar concentrations can cause up or downregulation of LamB^36^) or osmolarity (known to impact OmpF expression^37^). These non-genetic phenotypic shifts could contribute to environmental heterogeneity necessary to cause negative-frequency dependent selection on receptor tropism.

The second hypothesis suggests that rapid bacterial evolution, driven by mutations that downregulate one receptor and then the other^38^, could account for the observed fluctuations and emergence of negative-frequency dependent selection^39^. This alternative explanation focuses on genetic changes rather than phenotypic plasticity.

Both hypotheses offer plausible explanations for the observed gene expression shifts, and further investigation was required to determine which mechanism, or possibly a combination of both, is responsible for the phenomenon.

We focused on testing the evolutionary hypothesis through two separate studies conducted concurrently with the previously described temporal selection experiments. In the first trial, we sampled 1/10^th^ of bacterial populations for genomic DNA at two-hour intervals (0, 2, 4, 6, and 8 hours) across three replicates. Whole-genome sequencing revealed that a single mutation reached appreciable frequencies, a single amino acid change in the *envZ* gene, rising to low frequencies by 6 hours (10-18%) and dominating by the end (56-65%) in all three replicates (Fig. S2). Given the parallelism, this mutation likely preexisted in the bacterial culture the flasks were inoculated with. EnvZ is known to impact OmpF^40^ regulation and affect phage sensitivity^41^. While this mutation could potentially explain some of the reduction in OmpF expression at the conclusion of the 8-hour trials, it cannot account for the initial decrease. Furthermore, the unidirectional spread of the envZ mutation cannot explain the observed oscillations in LamB and OmpF expression throughout the trial.

In the second temporal selection experiment, we sampled three bacterial isolates from three replicate flasks every two hours and evaluated their sensitivity to λF and λL. Resistance was assessed by comparing the plaque-forming units of each phage on the isolates relative to the wild type (WT), a measurement known as efficiency of plaquing (EOP). A decline in EOP over time indicates an increase in resistance. Our results corroborated the genomic data: on average, *E. coli* exhibited a gradual evolution of resistance to λF, but not to λL (Fig. 5). This finding is consistent with the predicted phenotypic effect of the *envZ* mutation, which is associated with the repression of OmpF.

**Figure 5:**
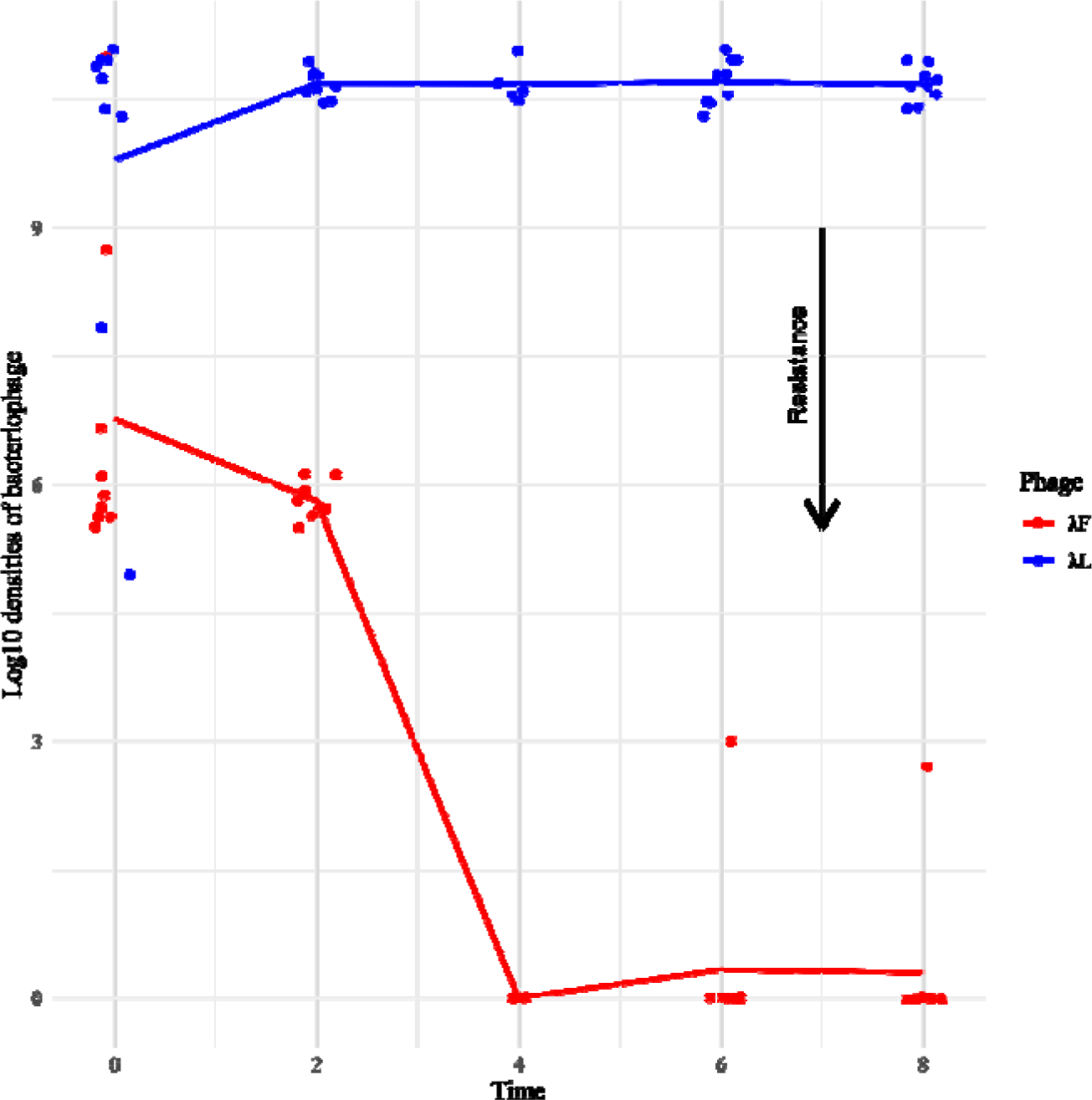
λF resistance evolves within an 8-hour period. Each point indicates how well either λF or λL can form plaques on a host isolated from different times and replicate flasks. More plaques means greater sensitivity to the phages. WT sensitivity is described by the values at x-axis ‘0’, λL starts out higher than λF because λL tends to have higher tighter cultures.

Our findings reveal the potential for significant evolutionary changes within mere hours, influencing not only species interactions but also the evolution and maintenance of biodiversity. However, the observed patterns—specifically, the lack of significant resistance to λL and the absence of cycling resistance patterns that mirror the oscillations in gene expression and λ genotypic frequency—suggest that rapid genetic evolution alone cannot fully explain the phenomena. The discrepancy between the unidirectional genetic changes and the oscillatory gene expression patterns strongly implies that non-genetic factors play a crucial role alongside evolutionary genetic mechanisms in generating the observed heterogeneity. While the relative contributions of genetic and non-genetic factors remain to be quantified, our studies collectively demonstrate that cryptic variation in receptor gene expression, arising from this interplay of mechanisms, can generate negative frequency-dependent selection and drive phage diversification. This underscores the complex and multifaceted nature of microbial adaptation and community dynamics.

### Conclusion

Phages are believed to possess the greatest diversity of life on Earth, challenging our understanding of species richness. Their extraordinary diversity, which surpasses that of their hosts, defies conventional ecological theories. Our study revealed phages’ remarkable propensity to diversify, even under controlled conditions designed to minimize diversification.

Despite our efforts to maintain isogenic bacterial cultures, phages found paths to diversification. This was due to inadvertent host heterogeneity introduced through temporal variability in gene expression and rapid evolution. Remarkably, this was sufficient to promote repeated diversification in less than 12 days (35 8-hour cycles) of evolution.

Our findings align with recent studies demonstrating the importance of rapid evolution in shaping phage ecology and diversity. For instance, rapid evolution of bacterial resistance to a generalist phage was shown to facilitate coexistence between the generalist and a specialist phage. Initially, rapid evolution was neither expected nor detected, and so, similar to our study, the maintenance of diversity seemed to contradict predictions based on competitive exclusion theory^42^. Both their observations and ours support theoretical models predicting that rapid evolution can promote coexistence and maintain biodiversity^43^.

Additionally, in line with our observations, cell-to-cell nongenetic variability plays a crucial role in maintaining phage species variation in another study. Pyenson et al. (2024) showed that different phage species adapt to cells in distinct growth phases, and that cells within populations naturally vary in their growth phases despite identical environments and genetic makeup. This heterogeneity alone can promote phage species coexistence^44^.

Efforts to minimize environmental heterogeneity were similarly foiled in a bacterial evolution experiment. Even in highly controlled chemostats with a single limiting resource, bacterial diversification has been observed^45^. This contradicts theoretical predictions favoring a single, most competitive genotype^46^. The discrepancy is attributed to tradeoffs not accounted for in the theory, such as between fast growth and stress tolerance resource^47^.

Collectively, these studies suggest that evolution by natural selection gravitates towards diversification more rapidly and resourcefully than often anticipated. Moreover, the conditions for coexistence are broader than expected, as organisms can partition their environments into increasingly finer niches. This understanding challenges and expands our current ecological theories, emphasizing the need for more comprehensive models that account for the complex interplay between rapid evolution, subtle environmental heterogeneity, and species coexistence.

## Methods

### Evolution experiment

To prepare the bacteria for each cycle, *Escherichia coli* strain BW25113 (WT)^48^ was grown exponentially in TrisLB media (10 g tryptone, 5 g yeast extract, 1 g (NH4)2SO4, 0.28 g K2HPO4, 0.08 g KH2PO4 per liter^38^. Autoclave, add 50 mL 1M Tris Base (pH 7.4), 200 µL CaCl2, and 10 mL MgSO4) at 37°C, 120 rpm for 2 hours to a density of ∼10^7^ cells/mL. On the first day, Bacteriophage λ strain EvoC (derived from λ reference genome GenBank: NC_001416^22^) was grown overnight to a density of ∼10^8^ virions/mL and filter sterilized.

The evolution experiment was initiated with six flasks containing 9.8 mL TrisLB media and were inoculated with 100 μL WT bacteria and 100 μL EvoC phage (MOI = 10). The flasks were incubated at 37°C, 120 rpm for 8 hours. At the end of the cycle, phage lysates were filtered through 0.22 µm filters and stored at 4°C. These lysates were then used to initiate the next cycle. This process was repeated on separate days for 35 cycles, with phage lysates stored at -80°C in 15% glycerol every 5th cycle.

Phages were sampled from lysates stored on cycles 25, 30, and 35 by plating on lawns of LamB^-^ (Keio collection JW3996) and OmpF^-^ (Keio collection JW0912)^48^ cells suspended in soft agar (10 g tryptone, 8 g NaCl, 1 g yeast extract, 1.75 g agar per liter. Autoclave, add 1.25 mL 20% glucose, 0.5 mL 1M CaCl2, and 2.5 mL 1M MgSO4 per 250 mL)^49^. Two individual phage clones were isolated from each plate for each replicate and then sequentially re-plated to guarantee a single clone was isolated. The specialization index of each phage sampled was calculated by plating on lawns of LamB^-^ or OmpF^-^ cells and measuring the density of plaque forming units on each host type. The index was calculated as: (density on OmpF^-^ - density on LamB^-^) / (density on OmpF^-^ + density on LamB^-^)^23^. The index ranges from -1 to +1 with -1 being a pure OmpF specialist and +1 indicating a perfect LamB specialist.

### Genetic Engineering of Blue-Marked Phage

We studied competition between a single OmpF- and LamB-specialist from population #2 of cycle 30. The phages were selected for their strong receptor preference and because they cooccurred in the same flask. To compete them head-to-head, we genetically marked each phage so that they would form blue plaques on a lawn of DH5α *E. coli* with 0.5 mg/ml 5-Bromo-4-Chloro-3-Indolyl β-D-Galactopyranoside (X-gal) and 0.25 mg/ml Isopropyl-beta-D-thiogalactoside (IPTG) supplemented^50^. We inserted a functional lacZα gene segment into their genomes via homologous recombination using *E. coli* cells containing a plasmid with the phage R gene fused to lacZα (provided by Ing-Nang Wang, SUNY Albany)^32^ and λ’s endogenous recombination system called λ-red^51^. Resulting phages were plated on a common *E. coli* strain that has lacZα deleted (DH5α) with X-gal and IPTG and blue plaques were chosen as marked specialists, and receptor preference was confirmed on LamB^-^ and OmpF^-^ plates.

### Frequency-dependent fitness measurements

Four receptor specialist phages (λF, λL, and their blue-marked versions) were grown overnight at 37°C and 120 rpm in 4 mL TrisLB with 100 μL LamB^-^ or OmpF^-^ and then filtered through 0.22 µm filters. Phage densities were estimated through counting plaque forming units on WT bacterial lawns. Mixed populations were prepared with 99% blue-marked phage and 1% unmarked phage. Six flasks with 9.8 mL TrisLB and 100 μL WT bacteria were inoculated with 100 μL of these mixtures, three with Blue-λL/Unmarked-λF and three with Blue-λF/Unmarked- λL. Cultures were grown at 37°C and 120 rpm for 8 hours. Phage lysates were filter-sterilized, and their densities were measured on overlay plates with DH5α, X-gal, and IPTG. Selection rates were calculated based on the changes in densities of blue and clear plaques. Selection rate is the difference of Malthusian growth rates divided by time^52^.

### Temporal selection studies

Twice we reconstituted population #2 at cycle 30 by setting up flasks identically to the original experiment, but this time using the two specialist phages that we genetically marked. We set up a total of 15 flasks and destructively sampled three of them every two hours (hours 0, 2, 4, 6, and 8). The flasks were setup with a mixture of λL unmarked and λF marked. On the first replay, phages were plated to determine their relative frequencies, and bacterial populations were sub-sampled for genome sequencing. In the second trial, phage frequencies were quantified, and bacteria were sampled to isolate strains to test their sensitivity to phage and the remaining population was used to extract RNA for gene expression analyses. Coincidentally, our initial phage mixtures were not perfectly balanced and in the first trial, the λL was more abundant and in the second the λF was more abundant. This led to a fortuitous observation of reciprocal oscillations.

### Illumina Sequencing and Breseq Analysis

1 mL bacterial samples from each flask from the first trial of the temporal selection experiment and were used to extract genomic DNA via Invitrogen Purelink Genomic DNA Kit. The DNA was sent for Illumina Whole Genome Sequencing to Seq Center, Pittsburgh PA. Mutations were identified using the breseq (version v0.38.0) computational pipeline in polymorphism mode^53^, referencing the BW25113 genome sequence (GenBank™ CP009273.1). Initially we ran breseq under standard polymorphism settings, but it identified hundreds of mutations spread throughout the full genome at low frequency, which were likely false positives. Although, one mutation in *envZ* was identified as reaching high frequencies. We repeated the breseq analysis, this time with more restrictive cutoffs, mutations were only identified if they were observed in 20 reads (average coverage was 180 reads). This second analysis removed the false positives and still identified the *envZ* mutation, an intergenic mutation (kgtP ← / ← rrfG; 321/+2), as well as hundreds of mutations in prophage genes, which are caused by cross contamination with λ genomic DNA since homologous phage genes occur in *E. coli*’s genome. Mutations in prophage genes were removed from the analyses.

### Bacterial Resistance Measurements

During the second one-cycle replay, bacterial colonies were streaked onto LB agar plates every two hours, incubated overnight, and re-streaked three more times to isolate bacteria away from phage. Single colonies were then grown overnight in TrisLB, and samples were stored in 15% glycerol. Phage resistance was measured by spotting serially diluted phage lysates on bacterial lawns and counting plaques^54^. WT control lawns were also spotted on to compare the number of plaques that form on a sensitive strain of bacteria.

### Phage Density Determination

Phage densities were determined using spot assays on WT lawns grown on overlay plates. Plates were incubated overnight at 37, plaques were counted, and densities as plaque-forming units per milliliter were calculated.

### Gene Expression Measurements Through Quantitative PCR

∼10 mL of bacterial samples from the second temporal replay experiment were removed, centrifuged, and pellets were resuspended in TRIzol reagent. RNA was extracted (Invitrogen TRIzol™ Plus RNA Purification Kit protocol with additional On–Column PureLink™ DNase Treatment), concentration measured using Thermo Scientific Nanodrop One spectrophotometer, and converted to cDNA using Applied Biosystems High-Capacity RNA-to-cDNA™ Kit. Gene expression of LamB, OmpF, and GapA (a constitutively expressed house-keeping gene) was analyzed using Real-Time PCR with SYBR Green. PCR solutions were created according to Bio Rad’s iTaq Universal SYBR Green Supermix using the 10uL reaction recipe for genomic content on a C1000 Touch Thermal Cycler machine with a CFX96 Touch Real-Time PCR Detection System attachment to perform the optical analysis. Standard curves and controls were included, and data was analyzed using Bio-Rad’s CFX Maestro software.

Before running the experiment, controls were run on WT, LamB^-^, and OmpF^-^ cells and gene expression profiles were in line with expectations (Fig. S3). A standard curve was generated to ensure consistent fluorescence in all runs. The standard was created through two replicates of an hour 0 bacterial sample (WT) that were serially diluted from 10^0^ to 10^-7^. Non-template controls as well as a housekeeping gene (*gapA*) were utilized for further validation of results and for calculating the final expression results.

### Primers for Real-Time PCR

Primers were designed and synthesized by Integrated DNA Technologies (Integrated DNA Technologies, San Diego location) with the following criteria: GC content of 40-60%, length of 18-30 bp, and amplicon size of 50-150 bp. Primers were free of runs of 4 or more base pairs, dinucleotide repeats, and ended with either a G or C nucleotide. BLAST checks ensured binding specificity to target sites of LamB, OmpF, and GapA genes. Although LamB-Reverse and OmpF-Forward primers showed some self-dimer possibilities, these were limited to 4 bp in length with ΔG values of -10.36 kcal/mol and -9.89 kcal/mol, respectively, which are just below the recommended -9 kcal/mol threshold for Real-Time PCR primers^55-58^. The primer sequences used were: LamB-Forward, 5’ TCG ATG TTG GCT TCG GTA AC; LamB-Reverse, 5’ TCG CGG TTT CGT TGG TAT AGT C; OmpF-Forward, 5’ GCG CAA TAC CAG TTC GAT TTT C; OmpF-Reverse, 5’ ACC AGA TCA ACA TCA CCG ATA C; GapA-Forward, 5’ CTG CTG AAG GCG AAA TGA AAG; and GapA-Reverse, 5’ TTC AGA GCG ATA CCA GCT TAG.

### Sanger Sequencing

The reactive region of gene *J* was sequenced using Sanger sequencing. PCRs were amplified using Forward 5’ CCT GCG GGC GGT TTT GTC ATT TA; Reverse 5’ CGC ATC GTT CAC CTC TCA CT ^59^. Only the reverse primer was used for sequencing, providing about 900 bases providing sequence of the full receptor-binding domain as well as other sections of the protein. Sequencing was performed by Azenta, La Jolla, CA, sequences were aligned using CLC Viewer 7^60^, and mutations called by visual inspection of the alignment.

## Acknowledgements

This work was supported by NSF grant 1934515 to JM. We acknowledge the helpful inputs of Josh Borin, Mike Doud, Krista Gerbino, Justin Lee, Matthew Daugherty, and Katherine Petrie. We also acknowledge Sweetzel Labador for laboratory assistance and Sahana Kuthyar for Real-Time PCR instruction.

## Supporting information

**Fig. S1:**
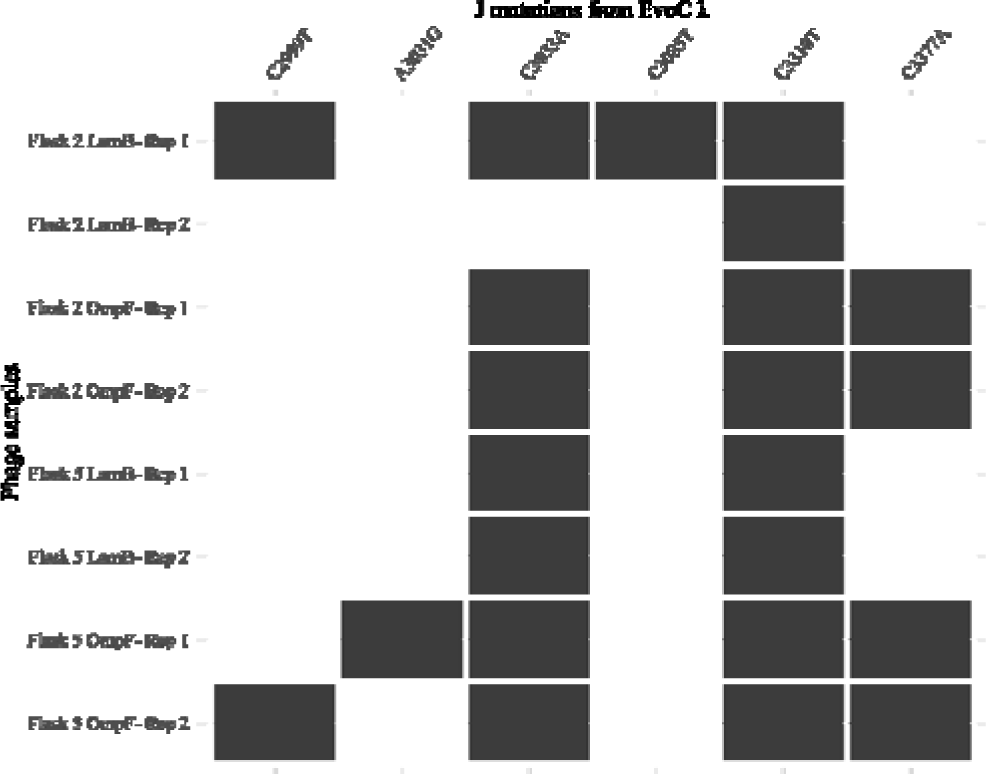
Receptor-binding domain mutations in J uncovered in isolates from populations #2 and #5 that were isolated using LamB^-^ or OmpF^-^ host lawns. 2 replicates per host and flask. All mutations are nonsynonymous and evolved in parallel in the two flasks except for A3031G and C3085T, which were both synonymous mutations.

**Fig. S2:**
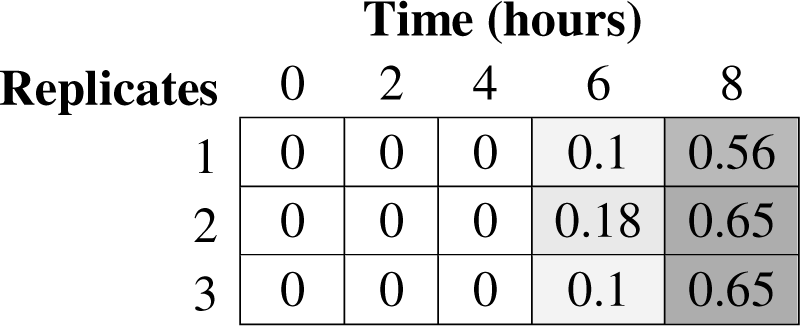
*envZ* mutation (L84R; CTG→CGG) rise in frequency over 8 hours.

**Figure S3:**
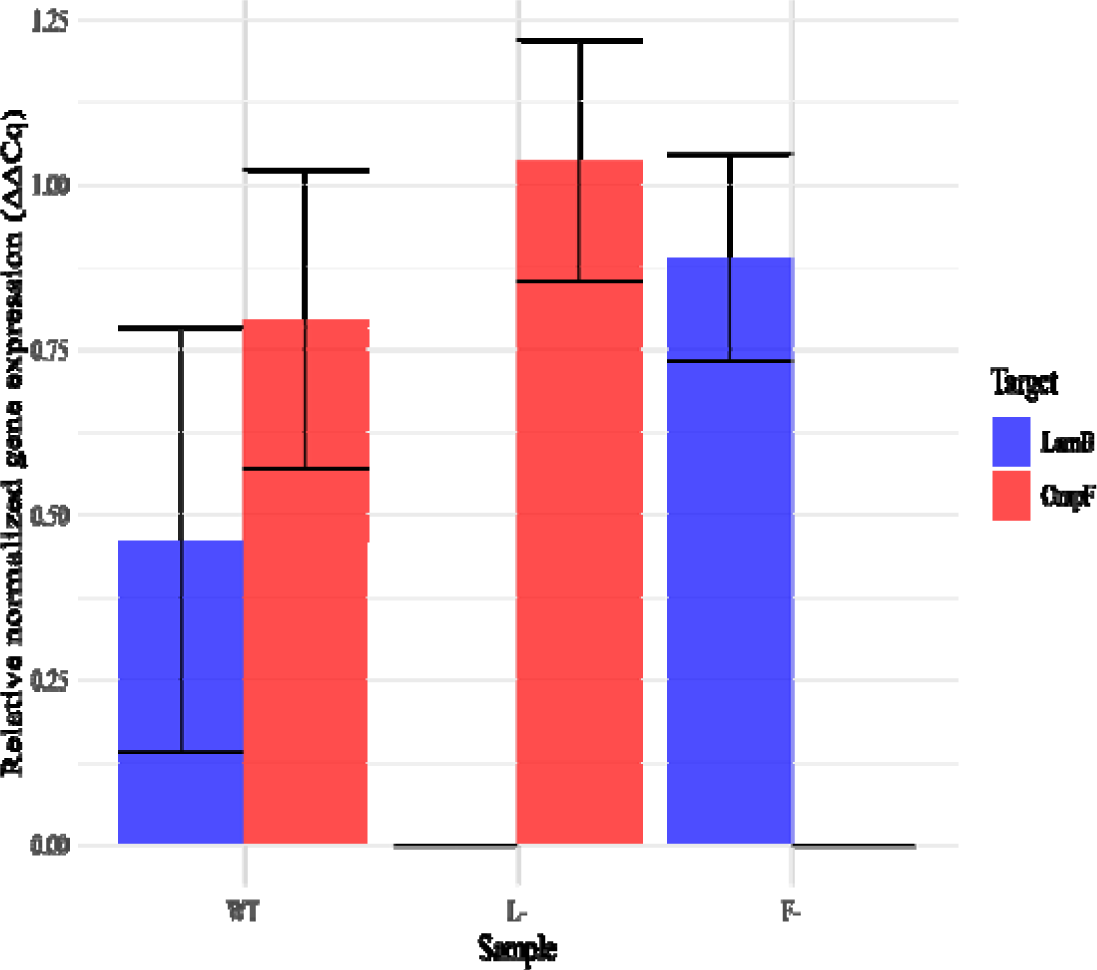
Gene expression controls for LamB and OmpF. Transcripts for an isolate expected to express both LamB and OmpF (WT) and those expected to only express one or the other receptor. Expression levels were normalized to the levels of a constitutively express housekeeping gene, *gapA*. Error bars represent standard deviations, with each bar indicating the average of triplicate measurements, except for F- in which an error resulted in only two replicates.

## Notes

### Competing Interest Statement

The authors have declared no competing interest.

